# Mycn Reactivates the Cell Cycle in Adult Cardiomyocytes and Promotes Cardioprotection in Myocardial Infarction

**DOI:** 10.1101/2025.07.23.666231

**Authors:** Aina Hirofuji, Hiroki Tanaka, Yuki Tsujita, Ryo Okubo, Ryohei Ushioda, Yumiko Fujii, Yusuke Ono, Michihiro Hashimoto, Megumi Kanao-Kanda, Yusuke Mizukami, Hiroyuki Kamiya, Kyohei Oyama

## Abstract

**Background:** Adult mammalian cardiomyocytes are terminally differentiated and possess minimal capacity for cell cycle re-entry. However, recent studies have shown that a rare population of cycling cardiomyocytes may confer cardioprotective effects following myocardial infarction (MI). The Myc family—Myc, Mycl, and Mycn—plays central roles in regulating cell cycle progression and cellular plasticity. This raises the possibility that specific Myc isoforms could reactivate the cardiomyocyte cell cycle and promote cardioprotective responses in the ischemic heart. This study aimed to systematically evaluate the effects of Myc, Mycl, and Mycn on cardiomyocyte cell cycle activation and cardiac outcomes following MI.

**Methods:** Cardiomyocyte-specific overexpression of *Myc*, *Mycl*, or *Mycn* was achieved in adult mice via adeno-associated virus 9 vectors driven by the cardiac troponin T promoter. Cardiomyocyte cell cycle activity, structural remodeling, and functional outcomes were assessed using RNA sequencing, immunohistochemistry, and transthoracic echocardiography. Cardioprotective effects were evaluated in MI model mice.

**Results:** While all three Myc family members activated cell cycle-related gene expression to varying degrees, Mycn elicited the most robust response. *Mycn* overexpression significantly enhanced cardiomyocyte cell cycle re-entry, as demonstrated by increased 5-bromo-2’-deoxyuridine incorporation and phosphorylation of histone H3. Mycn also induced hypertrophic growth, reflected by increased cardiomyocyte size and heart mass. Transcriptomic analyses revealed that Mycn uniquely upregulated genes associated with extracellular matrix remodeling and paracrine signaling, typically enriched in neonatal cardiomyocytes and previously linked to cardioprotection. Functionally, Mycn expression preserved cardiac contractility after MI, reduced infarct size, and increased capillary density in peri-infarct regions.

**Conclusions:** These findings identify Mycn as a functionally distinct member of the Myc family in the adult heart, capable of reactivating the cardiomyocyte cell cycle and promoting cardiac protection after ischemic injury. The cardioprotective effects appear to be mediated, in part, by the induction of a neonatal-like gene expression program supporting a stress-adaptive cardiac microenvironment.

## Introduction

Cardiomyocytes in adult mammals, including humans, exhibit an extremely limited capacity for cell division, rendering the heart one of the least regenerative organs in the body.^1,2^ Following myocardial injury—such as myocardial infarction (MI) or myocarditis—irreversible fibrosis typically ensues, leading to progressive and permanent cardiac dysfunction. In contrast, neonatal mammals retain an intrinsic capacity for cardiomyocyte proliferation during a narrow postnatal window—within one week of birth in mice—which enables them to regenerate heart tissue.^3,4^ This observation has spurred investigations into the molecular mechanisms governing cardiomyocytes cell cycle, with the aim of developing new strategies to support cardiac repair or protection. Postnatal mammalian cardiomyocytes are widely considered terminally differentiated and rarely re-enter the cell cycle. However, accumulating evidence suggests that a small subset of cardiomyocytes retains the ability to re-engage with the cell cycle at a very low frequency—approximately 0.1% per year in both humans and mice.^5,6^ This activity appears to be enhanced under pathological conditions; for example, in surgically induced MI model mice, cardiomyocytes in infarct border zones exhibit a marked increase in cell cycle re-entry.^7,8^ Notably, genetic ablation of endogenously cycling cardiomyocytes significantly worsens cardiac function following MI in mice.^9^ Although the mechanism remains to be fully elucidated, these findings suggest cardiomyocyte cell cycle activation confer cardiac protection in hearts under ischemic stress.

The Myc family of transcription factors—comprising Myc, Mycl, and Mycn—are well-established regulators of cellular proliferation, metabolism, and microenvironmental remodeling. Each member exhibits tissue- and context-specific functions;^10,11^ for example, *Mycl* is critical for pancreatic cell division,^12^ while *Mycn* plays a critical role in cardiomyocyte proliferation during embryonic heart development.^13,14^ Given these roles, the promotion of cell cycle activity is considered a core function of the Myc gene family. However, previous studies have shown that forced expression of *Myc* alone in adult cardiomyocytes induces only partial cell cycle re-entry and fails to drive full proliferation.^15,16^ In contrast, the functional roles of *Mycl* and *Mycn* in mature cardiomyocytes remain largely uncharacterized, raising the possibility that they may differ in their capacity to activate the cardiomyocyte cell cycle.

We hypothesized that individual Myc isoforms possess distinct functional capacities in adult cardiomyocytes, with specific isoforms capable of promoting more cell cycle activation and providing protection in the setting of MI. To test this hypothesis, we induced cardiomyocyte-specific overexpression of *Myc*, *Mycl*, and *Mycn* in adult mice and compared their effects on cell cycle activation. Additionally, we assessed whether any Myc family member conferred functional cardioprotection in the context of MI.

## Method

### Construction of AAV Transfer Vectors

A cTnT promoter-driven *GFP*-expressing adeno-associated virus (AAV) transfer plasmid (pENN.AAV.cTNT.PI.eGFP.WPRE.rBG; Addgene #105543) was used as the backbone. The plasmid was digested with BmtI and KpnI to replace the GFP coding sequence with *GFP*-V5, *Myc*-V5, *Mycl*-V5, or *Mycn*-V5 using the In-Fusion HD Cloning Kit (Clontech, 639649). The sequences of the inserted constructs are provided in **Supplemental File 1**. The resulting AAV transfer plasmids (pAAV-cTnT-*GFP*-V5, pAAV-cTnT-*Myc*-V5, pAAV-cTnT-*Mycl*-V5, and pAAV-cTnT-*Mycn*-V5) were packaged into AAV9 particles following a protocol described by Kimura and colleagues, ^17^ or using the AAV production service provided by VectorBuilder.

### Animal studies

All mice used in this study were housed under specific pathogen-free conditions at the Asahikawa Medical University. All animal experiments were approved by the Institutional Animal Care and Use Committee of the Asahikawa Medical University.

### Mice and AAV Vector Administration

Male C57BL/6 mice were purchased from Japan SLC, Inc. At six weeks of age, AAV vectors (1.5 × 10¹¹ viral genome copies in 100 μl per mouse) were administered via tail vein injection. Starting on the day of AAV vector administration, mice were provided with drinking water containing 5-bromo-2’-deoxyuridine (BrdU, 0.4 mg/mL) continuously for two weeks for end-point assay of the number of cardiomyocytes that had re-entered the S-phase. At the end of the treatment period, cardiac function was assessed by transthoracic echocardiography (Venu Go, GE HealthCare), and hearts were harvested for downstream analyses.

### MI Model Mice: Permanent Left Anterior Descending Artery (LAD) Ligation

Male C57BL/6 mice at six weeks of age were randomly allocated to the control (GFP) group and Mycn groups and then randomly assigned to either sham or MI surgery. The mice were initially anesthetized in an induction chamber with isoflurane (2% volume, air 500 mL/min) and subcutaneously injected with buprenorphine (0.05 μg/g). The mice were placed on a sterile heating plate (ThermoStar, Intellibio) set at 40 °C in the supine position with a face mask connected to an anesthesia system (MK-V100, Muromachi). Isoflurane was adjusted to provide a maintenance level (1.4-2.4% volume, air 500 mL/min) throughout the procedure. Anesthesia was monitored by observing the depth and rate of respiration, mucous membrane color, reflexes and tail pinch, and the overall appearance of muscle relaxation.

All surgical procedures were performed using a surgical loupe (Orascoptic) and a sterile technique. The surgical sites (neck and chest wall) were shaved and prepped with 10% iodine. A small midline cervical incision of 2 mm was made, and the skin, muscle, and adipose tissue covering the trachea were gently separated with blunt surgical instruments to expose the j thoracic cavity using blunt forceps until the left lung and pericardium were exposed. The retractors were repositioned onto the upper and lower ribs of the intercostal space to secure the surgical field of the heart.

For sham surgery mice, the thoracic cavity was kept open for 10 seconds, without further manipulation within the thoracic cavity, and the chest wall and skin were sutured closed. In MI model mice, the pericardium was bluntly dissected to expose the LAD. The LAD was then ligated as centrally as possible using a 7-0 prolene suture (Ethicon). LAD occlusion was verified by the blanching of the left ventricular muscle downstream of the ligation. The retractors were removed, and as much air as possible was pushed out of the thoracic cavity before the chest wall and skin were sutured with a 5-0 prolene suture (Ethicon). Mice received postoperative oral analgesia via meloxicam (Sawai) in drinking water (0.02 mg/mL, for approximately 5 mg/kg/day) for 5 days. The mortality rate associated with surgeries was approximately 11% for sham mice and 37% for MI model mice.

### Transthoracic Echocardiography on Mice

Anesthesia maintenance and monitoring were performed in the same manner as during the surgical procedures. Mice were initially anesthetized in an induction chamber with isoflurane (2% volume, air 500 mL/min), and on a heating plate (ThermoStar, Intellibio) set at 40℃, gently restrained in the supine position, with a face mask delivering isoflurane anesthesia (1.4-2.4% volume, air 500 mL/min) throughout the echocardiography.

All echocardiography was performed with the L10-22 linear transducer probe and Venu Go echocardiography system (GE HealthCare), which included an integrated limb lead electrocardiogram (ECG). The ECG electrodes were secured topically on the limbs, the fur was gently removed with hair removal cream, and ultrasound contact gel was applied to the transducer probe. Initial B-mode images were acquired in the parasternal long-axis view, ensuring clear visualization of both the apex and aortic valves. Subsequently, B-mode short-axis images were obtained by rotating the ultrasound transducer approximately 90°, enabling the identification of the mitral valve papillary muscles, followed by M-mode images in the same visual field. Left ventricular dimensions were calculated from parasternal long-axis images using the Venu Go software.

### Cardiomyocyte Isolation

Cardiomyocytes were isolated using Langendorff perfusion digestion, as previously described, with minor modifications.^18^ Briefly, the hearts were washed via coronary perfusion with calcium-free Tyrode’s buffer (126 mM NaCl, 5.4 mM KCl, 0.33 mM NaH₂PO₄, 1 mM MgCl₂, 10 mM HEPES, 10 mM glucose, 20 mM taurine, pH 7.4) supplemented with 20 mM 2,3-butanedione monoxime (BDM; Cayman Chemical, 20828) for 2 min. Enzymatic digestion was then performed at 37 °C by perfusing with 0.69 U/mL Liberase TH (Roche, 5401151001) prepared in calcium-free Tyrode’s buffer containing 20 mM BDM for 10–12 min. The digested hearts were mechanically dissociated in ice-cold KB solution (20 mM KCl, 10 mM KH₂PO₄, 70 mM potassium glutamate, 1 mM MgCl₂, 25 mM glucose, 20 mM taurine, 0.5 mM EGTA, 10 mM HEPES, 0.1% albumin, pH 7.4). The resulting cardiomyocyte suspensions were passed through a 100–200 mm cell strainer to remove tissue debris and purified by low-speed centrifugation (100× g for 1 min), repeated three times. This procedure yielded cardiomyocytes with an approximate purity of 90%.

### RT-qPCR and RNA-seq

Total RNA was extracted from purified cardiac myocytes using TRIzol reagent (Thermo Fisher Scientific, 15596026) and further purified using the NucleoSpin® RNA kit (MACHEREY-NAGEL, 740955.250). Complementary DNA (cDNA) was synthesized from purified RNA using ReverTra Ace® qPCR RT Master Mix (TOYOBO, FSQ-201). Quantitative PCR (qPCR) was performed using the THUNDERBIRD® Next SYBR™ qPCR Mix (TOYOBO, QPX-201X5), according to the manufacturer’s instructions. The gene-specific primers used in this study are listed in **Supplemental File 2**.

RNA sequencing was performed by Rhelixa Inc. (Tokyo, Japan) using ribosomal RNA depletion and strand-specific library preparation methods. Paired-end sequencing (150 bp ×2) was performed, generating approximately 40 million reads for each sample. Each group included two biological replicates each. RNA-seq data were deposited in the DDBJ BioProject database under the accession number PRJDB35457.

### RNA-seq Data Analysis

Raw sequencing reads were preprocessed using fastp (v0.23.2) with default settings.^19^ Transcript abundance was quantified using Salmon (v1.9.0) with the --gcBias option enabled. ^20^ The reference transcriptome and corresponding GTF annotation files were obtained from GENCODE Release M25 (GRCm38.p6). Transcript-level abundances were summarized to the gene level using the tximport package (v1.30.0), and differential gene expression analysis was performed using DESeq2.^21^ Differentially expressed genes (DEGs) were defined as those with a false discovery rate (FDR) <0.05 and fold change ≥2.

A list of DEGs was subjected to Gene Ontology (GO) enrichment analysis using the Gene Ontology Consortium’s web-based enrichment tool (https://geneontology. org/). GO terms in the Biological Process category with FDRs <0.05 were considered significantly enriched. To reduce redundancy, enriched GO terms were further processed using REVIGO (http://revigo.irb.hr) with the default semantic similarity measure and a medium similarity cutoff (0.7). GO terms with a dispensability score <0.05 were retained as representative enriched terms.

### Histopathological Analysis

For histopathological analysis, formalin-fixed paraffin-embedded (FFPE) tissue specimens were prepared. For bright-field microscopy, sections were stained using standard protocols for hematoxylin and eosin (HE) and Picro-Sirius Red. For immunohistochemical analysis, sections underwent sequential treatment including deparaffinization, rehydration, quenching of endogenous peroxidase activity, and antigen retrieval. Sections were then incubated with the following primary antibodies: anti-V5 (Sigma-Aldrich, V8012), anti-BrdU (ThermoFisher Scientific, B35128), and anti-phospho-Histone H3 (Abcam, ab5176). Detection was performed using horseradish peroxidase (HRP)-conjugated anti-mouse and anti-rabbit secondary antibodies (Vector Laboratories). For immunofluorescent staining, sections were incubated with the following primary antibodies: anti-HSP47 (Abcam, ab109117) and anti-Ki67 (ThermoFisher Scientific, 740008T). Detection was performed using an anti-rabbit secondary antibody conjugated with Alexa Fluor 555 (ThermoFisher Scientific, A-31572) and an anti-rat secondary antibody conjugated with Alexa Fluor 647 (Abcam, ab150155). Cell membranes were stained using wheat germ agglutinin (WGA) conjugated with Alexa Fluor 488, and Hoechst 33342 (FUJIFILM, 346-07951) was used for nuclear counterstaining. Fluorescent and bright-field images were obtained using a BZ-X800 fluorescence microscope (Keyence Co.) and a NanoZoomer S60 virtual slide scanner (Hamamatsu Photonics Co.), respectively.

Mouse heart tissues obtained from MI and sham surgeries were fixed and transversely sectioned into 2-mm slices. Among these, the slice exhibiting the largest infarct area was selected for quantitative analysis. Virtual slides stained with Picro-Sirius Red, as well as immunohistochemistry for BrdU and phospho-histone H3, were analyzed using QuPath software to quantify infarct area and the percentage of positively stained nuclei.

### Statistics

Statistical analyses were performed using GraphPad Prism software. One-way analysis of variance (ANOVA) followed by Tukey’s post hoc analysis was performed for multiple comparisons, and unpaired two-tailed Student’s t-tests were used for two-group comparisons. Statistical significance was set at *p*<.05. Data visualization was performed using GraphPad Prism software and the ggplot2 package in R. Data are represented with biological replicates as individual points and bars as mean±SD.

## Results

### AAV-Induced Overexpression of Myc Isoforms in Adult Mouse Cardiomyocytes

To investigate the effects of Myc family factors on the cell cycle in adult murine cardiomyocytes, an AAV9 gene delivery system was used. AAV9 vectors encoding cardiac-specific *Myc*, *Mycl*, *Mycn*, and a *GFP* control were intravenously injected into 6-week-old mice, and the hearts were harvested for analysis two weeks post-injection (**Figure 1A**). RNA was extracted from purified cardiomyocytes, and the expression of *Myc*, *Mycl*, *Mycn*, and *GFP* was measured using real-time RT-PCR, confirming the overexpression of the Myc family factors (**Figure 1B**). Immunohistochemistry with a V5 tag-specific antibody demonstrated that each Myc family factor was specifically expressed in cardiomyocytes within the myocardial tissue (**Figure 1C**). These data confirmed the achievement of cardiac-specific expression of all Myc family members.

**Figure 1.**
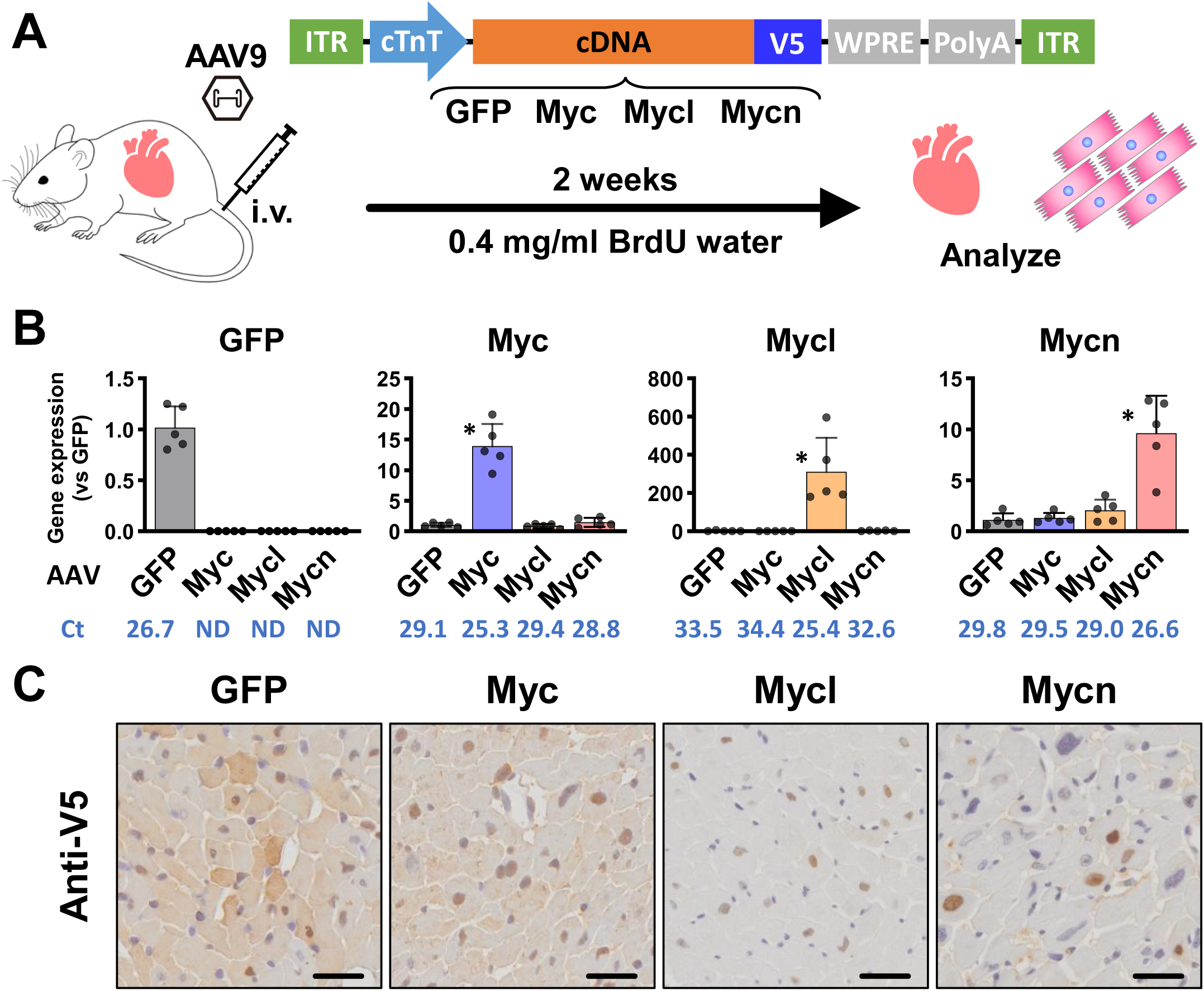
Cardiomyocyte-Specific Myc Family Gene Transduction Model. **(A) Experimental design:** Adeno-associated virus (AAV) vectors encoding *GFP*, *Myc*, *Mycl*, or *Mycn* were administered intravenously. Hearts or isolated cardiomyocytes were harvested two weeks post-injection for analysis. 5-bromo-2’-deoxyuridine (BrdU) was administered via drinking water during the two-week period for histological assessment of DNA synthesis. **(B) Validation of transgene expression:** Cardiomyocytes were isolated, and real-time RT-PCR was used to confirm expression of Myc family genes. \**p*<.01 vs. GFP. **(C) Validation of Cardiomyocyte-specific expression:** Immunostaining confirmed cardiomyocyte-specific expression of the transgenes. Scale bar: 25 µm.

### The Effects of Myc Family Factors on the Transcriptome in Cardiomyocytes

To determine whether there are functional differences among Myc family members, RNA sequencing was performed on purified cardiomyocytes two weeks after AAV9-mediated Myc family gene delivery. Differentially expressed gene (DEG) analysis revealed that *Myc*, *Mycl*, and *Mycn* upregulated 102, 25, and 367 genes and downregulated 20, 10, and 164 genes, respectively (**Figure 2A**). The Venn diagram indicated a strong overlap in upregulated genes, with approximately 80% of the upregulated genes in the Myc and Mycl groups included in the upregulated genes in the Mycn group (**Figure 2B**). In contrast, although the total number of downregulated genes in the Myc group was smaller than in the Mycn group, 70% of the downregulated genes in the Myc group did not overlap with those in the Mycn group. GO enrichment analysis revealed that all Myc family members upregulated genes related to cell cycle regulation, with lower FDR and higher gene numbers in the Mycn group compared to the Myc and Mycl groups. This includes GO terms such as “cell division (GO:0051301)” and “cell cycle (GO:0007049)”. Additionally, the Mycn group showed unique GO term enrichment, such as “collagen metabolic process (GO:0032963)” (**Figure 2C**). These data indicate that while all Myc family members commonly activate cell cycle-related genes, *Mycn* has the most significant impact and unique effects, particularly influencing the reorganization of the surrounding tissue environment, including cell adhesion and collagen production. Consistent with the RNA-seq data, RT-qPCR demonstrated that *Mycn* significantly upregulated G2/M and mitotic cell cycle regulator genes (*Ccnb1* and *Plk1*) and collagen genes (*Col1a1* and *Col1a2*) (**Figure 2D**).

**Figure 2.**
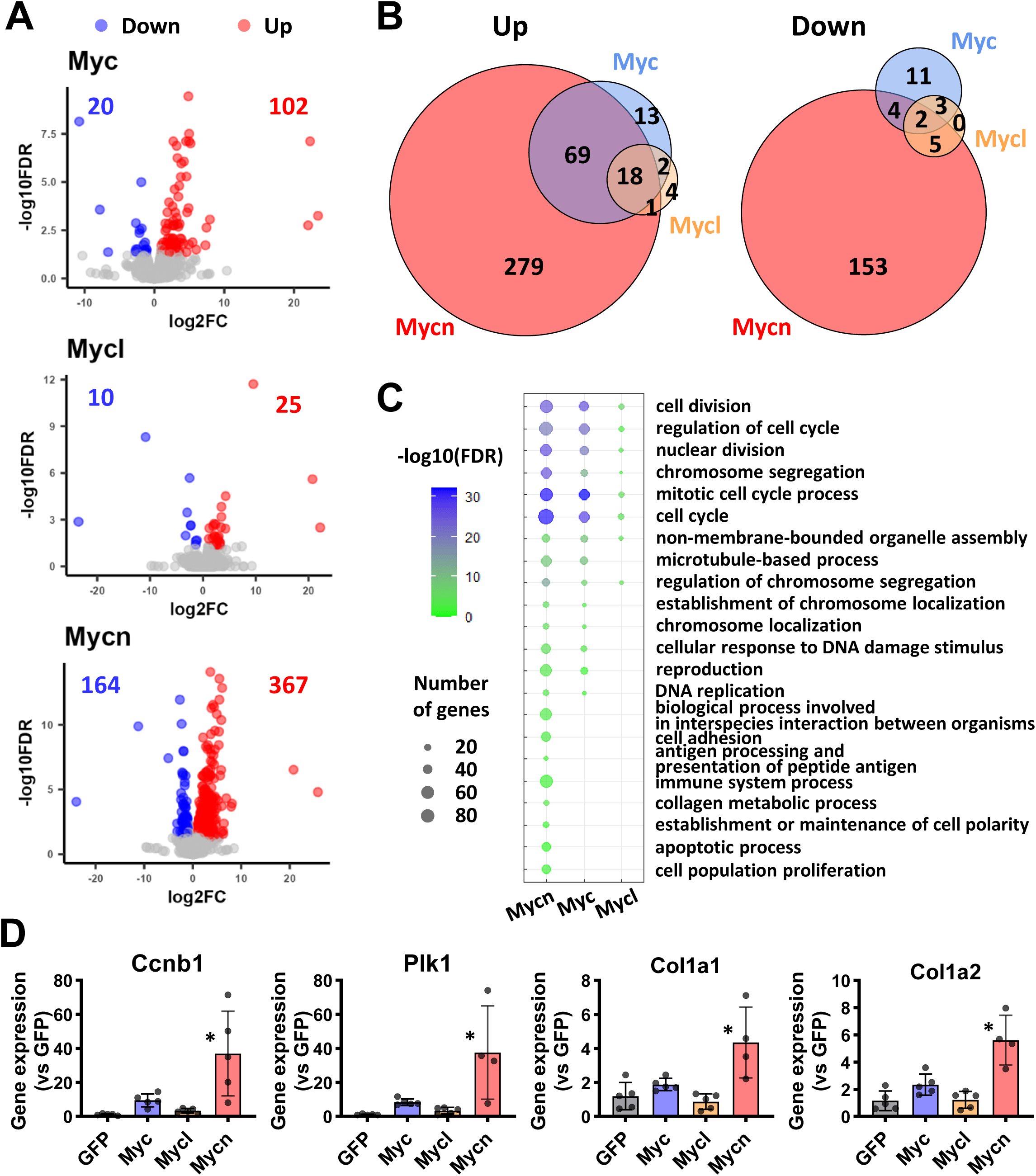
Transcriptional Impact of Myc Family Gene Expression in Cardiomyocytes. **(A) Differential gene expression:** RNA-seq was used to identify differentially expressed genes relative to the GFP control group. Genes were defined as differentially expressed with false discovery rate (FDR) <0.05 and an absolute fold change (FC) ≥2. Numbers of upregulated and downregulated genes are shown in red and blue, respectively. **(B) Venn diagram:** The numbers of commonly and uniquely altered genes among the groups are indicated. **(C) Gene Ontology (GO) enrichment analysis:** Enriched GO terms in biological processes for upregulated genes are shown. **(D) Validation of RNA-seq results:** Quantitative RT-PCR was performed for selected genes involved in G2/M phase progression and collagen metabolism. Dots represent individual biological replicates. \**p*<.05 vs. GFP.

### Mycn Induces Mitosis and Hypertrophy in Cardiomyocytes

As gene expression analysis showed that all Myc family members upregulated cell cycle genes, we next examined whether Myc family members affected cell cycle activity and heart growth. To label cardiomyocytes that had entered the S phase, BrdU, a thymidine analog, was administered via drinking water for two weeks after AAV delivery (**Figure 1A**). BrdU-positive cardiomyocytes were undetectable or barely detectable in the GFP, Myc and Mycl groups, the Mycn groups convincingly showed BrdU-positive cardiomyocytes, with the highest rate of BrdU-positive nuclei observed in the Mycn group heart tissue (2.3±2.0% in Mycn, 0.23±0.26% in GFP, 0.75±1.0% in Myc and 0.25±0.23% in Mycl) (**Figures 3A, 3B** top). Additionally, the mitotic marker phospho-H3 (Ser10) (pH3)-positive cardiomyocytes were not detected in neither the Myc nor Mycl groups; only the Mycn group showed convincing pH3-positive cardiomyocytes, with the highest rate of pH3-positive nuclei in the Mycn group (3.8±1.1% in Mycn, 0.16±0.06% in GFP, 0.29±0.04% in Myc and 0.55±0.38% in Mycl) (**Figures 3A**, **3B** bottom).

**Figure 3.**
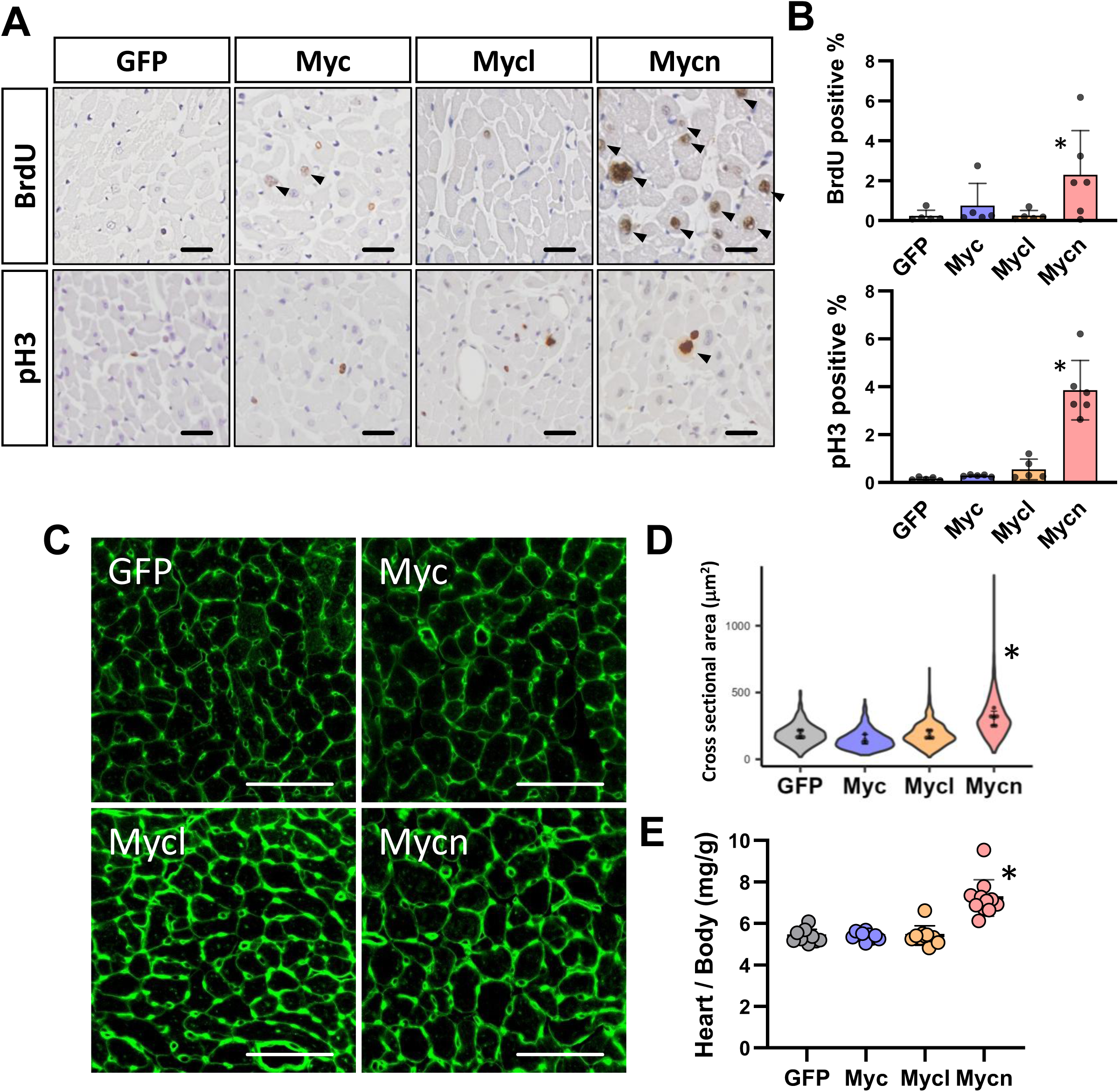
Myc Family Gene Effects on Cardiomyocyte Cell Cycle Activity and Cardiac Growth. **(A) Immunostaining for cell cycle markers:** 5-bromo-2’-deoxyuridine (BrdU) and phospho-Histone H3 (pH3) immunostaining were performed two weeks after adeno-associated virus (AAV) (*GFP*, *Myc*, *Mycl*, *Mycn*) vector administration. Arrowheads indicate cardiomyocyte nuclei positive for BrdU or pH3. Scale bar: 50 µm. **(B) Quantification of cycling nuclei:** The percentages of BrdU^+^ and pH3^+^ nuclei were quantified using QuPath software. \**p*<.05 vs. GFP. **(C) Representative images of cardiac cross-sections:** Representative wheat germ agglutinin (WGA)-stained sections used to assess cardiomyocyte cross-sectional area. Scale bar: 50 µm. **(D) Quantification of cardiomyocyte cross-sectional area:** Cross-sectional areas of 100 cardiomyocytes per mouse were measured. Dots represent average values for biological replicates; violin plots depict distribution; error bars indicate SD. \**p*<.05 vs. GFP. **(E) Cardiac Growth:** Heart weight normalized to body weight was assessed. Dots represent individual biological replicates. \**p*<.05 vs. GFP.

Cell cycle activation is associated with increased cell size and proliferation. To measure the cardiomyocyte size, wheat germ agglutinin (WGA) staining was performed (**Figures 3C**, **3D**). The cross-sectional area of cardiomyocytes in the GFP group was 188±79 µm², with no significant difference observed in the Myc (149±72 µm²) and Mycl (185±84 µm²) groups. In contrast, cardiomyocyte cell size in the Mycn group (307±141 µm²) was significantly larger than in the GFP, Myc, and Mycl groups (**Figure 3D**). Consistent with the increase in cell cycle activity and cell size, heart mass normalized to body weight was significantly increased in the Mycn group (7.2±0.84 mg/g) compared to the GFP, Myc, and Mycl groups (5.4±0.30 mg/g, 5.40±0.18 mg/g, 5.41±0.45 mg/g, respectively), likely due to hypertrophic growth (**Figure 3E**). Thus, Mycn, but not Myc or Mycl, has the greatest impact on cell cycle re-entry, inducing mitosis and hypertrophy in cardiomyocytes.

### Cardiac-specific Mycn Expression Activates Fibroblasts while Preserving Cardiac Function

We next assessed whether there were any pathophysiological changes in the hearts expressing Myc family members. Gene expression analysis indicated that *Mycn* upregulated genes related to the collagen metabolic process, in addition to cell cycle regulatory genes (**Figure 2**). Therefore, we first investigated whether the expression of cardiac-specific Myc family members alters fibroblast activation. Hsp47 (Heat shock protein 47) is a chaperone in collagen metabolism and a marker of myofibroblast activation.^22^ Immunofluorescence staining showed a significant increase in Hsp47-positive fibroblasts in *Mycn*-expressing heart sections (0.55±0.053 cells/mm²) compared to all other groups (GFP: 0.11±0.024 cells/mm², Myc: 0.22±0.043 cells/mm², Mycl: 0.26±0.031 cells/mm²) (**Figures 4A**, **4B**). Similarly, the number of fibroblasts positive for Ki67, a proliferation marker, was significantly increased in the Mycn group (GFP: 0.059±0.012 cells/mm², Myc: 0.20±0.073 cells/mm², Mycl: 0.17±0.040 cells/mm², Mycn: 0.32±0.054 cells/mm²). Consistent with these data, Picro-Sirius Red staining demonstrated a significant increase in interstitial collagen fibers, functioning as a structural framework between cardiomyocytes, in the Mycn group relative to the other groups (GFP: 0.22±0.038%, Myc: 0.16±0.054%, Mycl: 0.13±0.035%, Mycn: 0.63±0.44%) (**Figure 4C** top panel and **Figure 4D**). Some noticeable morphological changes were observed in the HE stained images. An increase in nuclear size and a speckled pattern of heterochromatic foci were observed in *Mycn*-expressing cardiomyocytes (**Figure 4C**, lower panel), which is probably related to cell cycle activation. Next, qPCR was performed to assess fetal gene expression, which is often related to pathological changes and activation of the cell cycle in cardiomyocyte.^23,24^ *Myh7* (Mycn: 6.2±2.7 fold, GFP: 1.1±0.6 fold, Myc: 1.2±0.42 fold, Mycl: 0.85±0.16 fold) and *Nppa* (Mycn: 4.2±2.3 fold, GFP: 1.1±0.4 fold, Myc: 0.82±0.4 fold, Mycl: 0.55±0.38 fold) gene expressions were significantly upregulated in the Mycn group compared to the other groups (**Figure 4E**). Finally, echocardiograms were performed to determine if the Myc family affect cardiac function, focusing on Myc and Mycn. As indicated in **Figure 4F**, neither *Myc* nor *Mycn* changed the ejection fraction up to 8 weeks after gene transduction, and no differences were observed compared to the GFP control group. These data indicate that although Mycn changes fibroblast activity and cardiac fetal gene expression, *Mycn*-expressing hearts exhibit normal heart function up to 8 weeks post-transgene induction.

**Figure 4.**
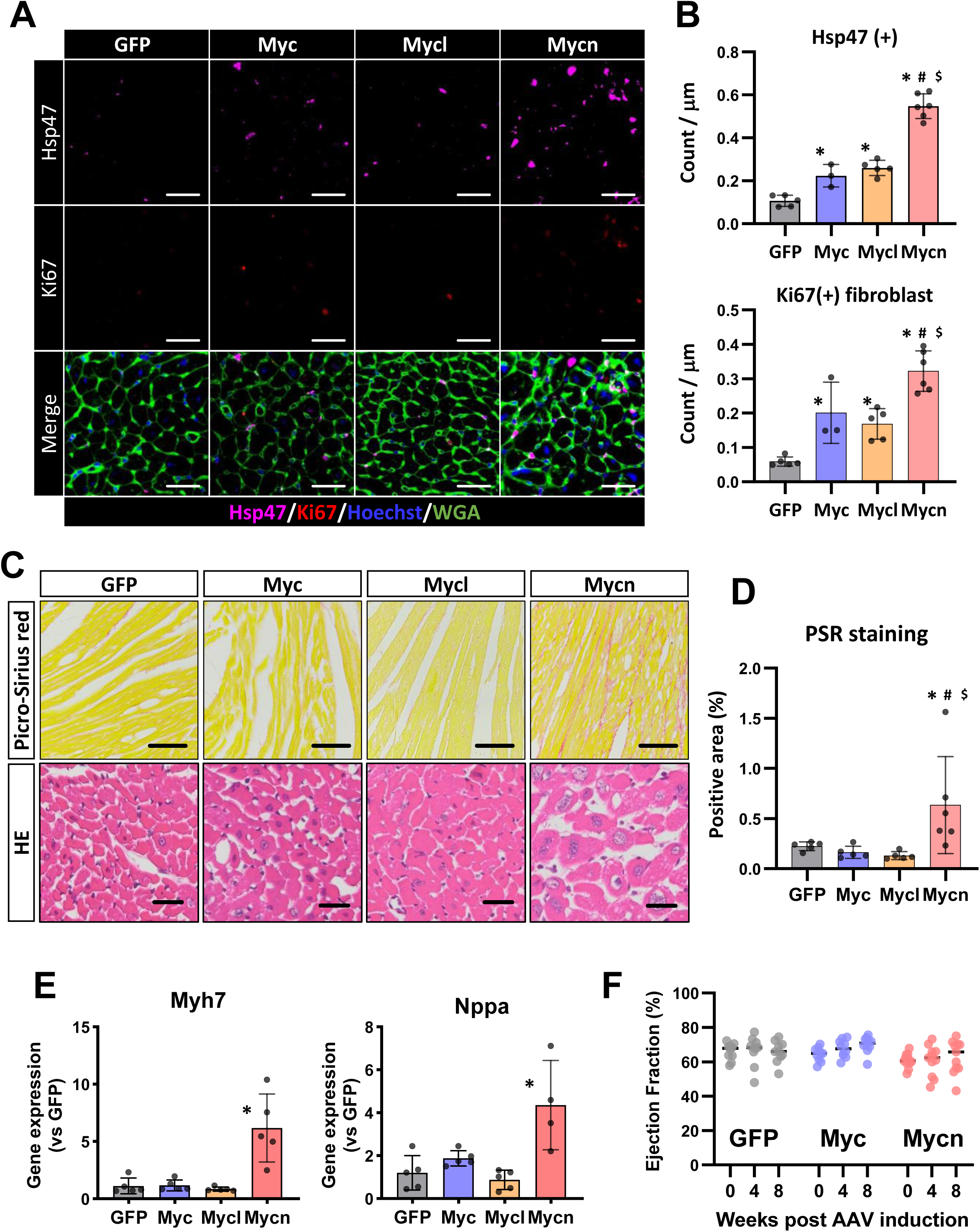
Myc Family Gene Effects on Fibroblast Activation and Cardiac Phenotype. **(A) Fibroblast activation markers:** Immunostaining for Hsp47 (magenta) and Ki67 (red) was performed to assess fibroblast activation two weeks after adeno-associated virus (AAV: *GFP*, *Myc*, *Mycl*, or *Mycn*) administration. Wheat germ agglutinin (WGA, green) was used to delineate cardiomyocytes. Scale bar: 30 µm. **(B) Quantification of fibroblast activation markers:** Numbers of Hsp47^+^ (top) and Ki67^+^ (bottom) fibroblasts per myocardial area were quantified. \**p*<.05 vs. GFP, ^#^*p*<.05 vs. Myc, ^$^*p*<.05 vs. Mycl. **(C) Histological assessment:** Representative images of Picro-Sirius Red (PSR, top) and hematoxylin-eosin (HE, bottom) staining. Scale bars: 100 µm (PSR) and 25 µm (HE). **(D) Quantification of collagen production:** PSR^+^ areas were quantified using QuPath software. \**p*<.05 vs. GFP, ^#^*p*<.05 vs. Myc, ^$^*p*<.05 vs. Mycl. **(E) Fetal gene expression:** RT-qPCR analysis of *Myh7* and *Nppa* expression. \**p*<.05 vs. GFP. **(F) Cardiac function:** Ejection fraction (EF) was assessed by transthoracic echocardiography at 0, 4, and 8 weeks following AAV transduction (*GFP*, *Myc*, *Mycn*).

### Mycn Preserves Heart Function after MI

Cycling cardiomyocytes have been reported to play a cardioprotective role.^9^ In contrast, cardiac fibroblasts can exert either cardioprotective effects by supporting tissue repair or contribute to adverse remodeling and progression of heart failure, depending on the context. Given that *Mycn* expression in cardiomyocytes had the greatest biological impact on heart tissue, we conducted further experiments to investigate the effects of *Mycn* expression on the injured myocardium. To this end, an MI model was employed on mice, which was induced via surgical ligation of the LAD. One day before MI surgery, echocardiography was performed to evaluate basal cardiac function, and AAV9 vectors (*GFP* or *Mycn*) were administered (**Figure 5A**). Four weeks later, cardiac function was measured, and the hearts were collected for histological analysis. Histological evaluation at 4 weeks after MI revealed extensive scar formation spanning the full thickness of the left ventricular wall in the *GFP*-expressing control group. In contrast, the *Mycn*-expressing group exhibited only a small, superficial scar confined to the subepicardial region (**Figures 5B-D**). Notably, the myocardium in the *Mycn*-expressing hearts showed increased vascular density in the peri-scar area, suggesting the formation of collateral vessels (**Figure 5B**). In the GFP group, consistent with the large scar formation, ejection fraction was significantly decreased at 4 weeks after MI compared to before MI (ΔEF −21.6±7.1%). In contrast, ejection fraction was maintained in the Mycn group after MI (**Figure 5E**). These data indicate that cardiac-specific *Mycn* expression prevented the decline in heart function after MI.

**Figure 5.**
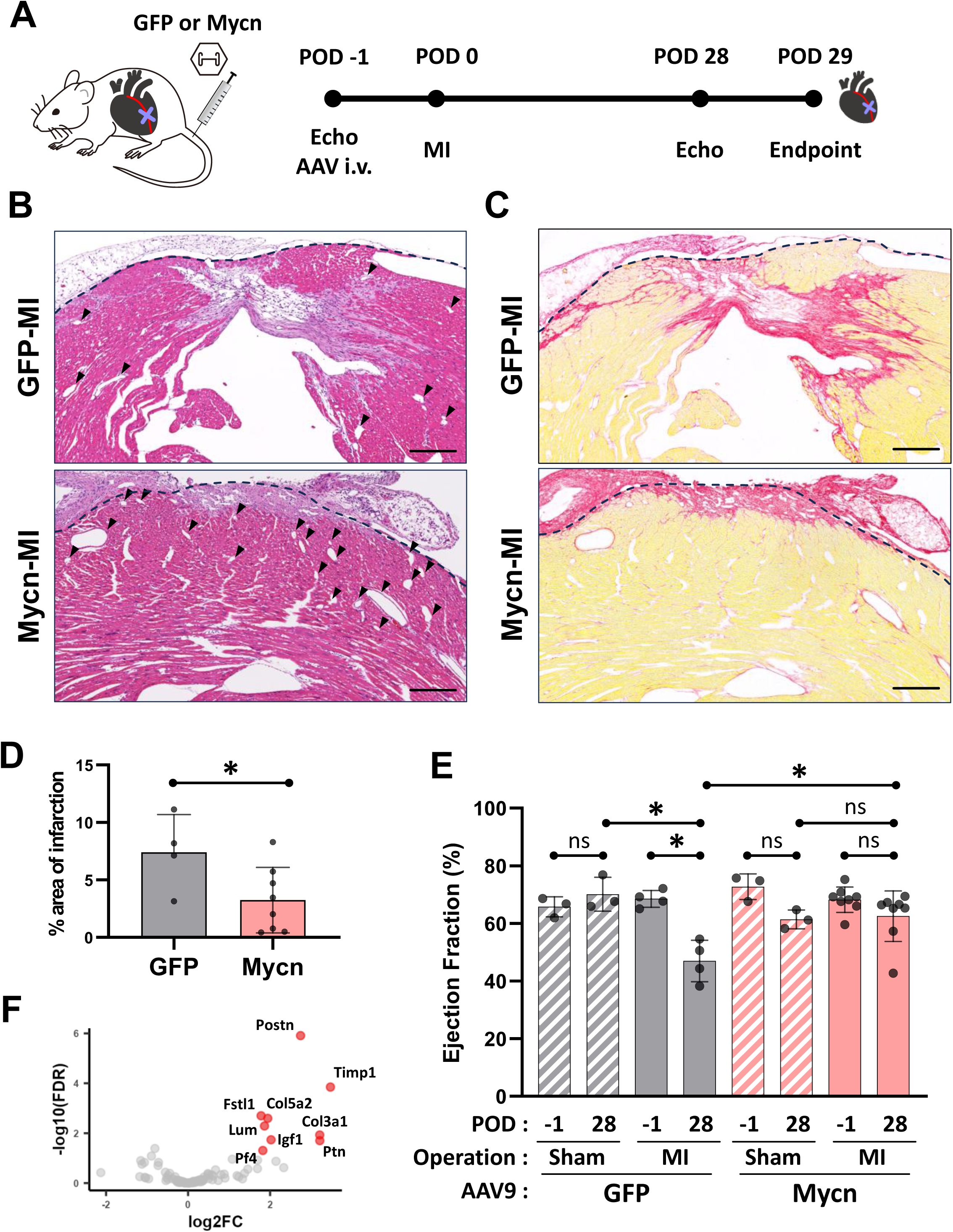
Mycn Preserves Cardiac Function after Myocardial Infarction (MI) **(A) Experimental Design:** Transthoracic echocardiography and adeno-associated virus (AAV: *GFP* or *Mycn*) vector administration were performed one day prior to MI surgery. Post-MI transthoracic echocardiography was performed on day 28, with histological analyses conducted on day 29. **(B, C) Representative images of post-MI histology:** Hematoxylin-eosin (HE, B) and Picro-Sirius Red (PSR, C) staining show myocardial architecture 28 days post-MI. Dashed lines indicate myocardial borders; arrowheads indicate vasculatures in (B). Scale bars: 250 μm. **(D) Quantification of fibrosis**: Fibrotic areas were measured using QuPath software. \**p*<.05 vs. GFP. **(E) Cardiac function after MI:** Ejection fraction (EF) was measured by thoracic echocardiography. Dots represent individual data points. \**p*<.05. **(F) Growth factor and angiogenic gene expression:** Differentially expressed gene (DEG) analysis was performed using RNA-seq data. Each dot represents a growth or angiogenic factor. Genes with false discovery rate (FDR) <0.05 and an absolute fold change (FC) ≥2 are shown in red.

Next, we explored the molecular mechanisms of cardiac protection by Mycn. The data showing a higher density of vasculature after MI (**Figure 5B**) and fibroblast activation in heart tissue (**Figures 4A-D**) by cardiac-specific *Mycn* expression suggested that changes in the cardiac microenvironment caused cardiac protection. Therefore, to find factors affecting the cardiac microenvironment, RNA-seq data was reanalyzed focusing on 95 genes having GO terms of “growth factor activity (GO:0008083)” or in the angiogenesis gene set.^25^ Of these, nine genes are significantly upregulated in the Mycn group. Strikingly, several genes have been identified to be involved in cardioprotection (*Fstl1*, *Timp1*, *Igf1*), angiogenesis (*Fstl1*, *Igf1*, *Ptn*), and fibroblast activation (*Postn*, *Col5a2*, *Col3a1*).^26–32^ These data suggest that Mycn-expressing cardiomyocytes modulated the cardiac microenvironment to resist irreversible cardiac injury, such as cardiomyocyte necrosis and myocardial tissue scarring.

## Discussion

In this study, we systematically investigated the functional effects of Myc family genes—*Myc*, *Mycl*, and *Mycn*—in adult murine cardiomyocytes using AAV-mediated cardiomyocyte-specific gene delivery. Our data reveal that among these paralogs, *Mycn* exerts the most distinct and pronounced biological effects, particularly in promoting cardiomyocyte cell cycle activation and modulating the cardiac microenvironment, such as fibroblast responses. Furthermore, in a murine MI model, *Mycn* expression in cardiomyocytes preserved systolic function and significantly reduced infarct area, suggesting a potent cardioprotective role. To our knowledge, this is the first study to directly compare the functional effects of the Myc family genes in the postnatal mammalian heart under ischemic stress, specifically MI. Our findings position Mycn as a potent modulator for enhancing cardioprotection through cardiomyocyte response and stromal remodeling during ischemic cardiac stress.

### Functional Divergence of Myc Family Members in Cardiomyocytes

Although the Myc family members share highly conserved basic helix-loop-helix-leucine zipper domains, they are not functionally redundant across tissues and developmental contexts.^11,12^ Our findings recapitulate this principle in postnatal cardiomyocytes, where we observed clear functional divergence among *Myc*, *Mycl*, and *Mycn*. Prior studies have shown that *Myc* overexpression in adult cardiomyocytes induces cardiac hypertrophy and DNA synthesis, but fails to elicit full mitotic re-entry, limiting its regenerative potential.^15,16,33,34^ Our findings align with these observations, as *Myc* promoted S-phase re-entry, but did not induce mitosis as indicated by the absence of pH3-positive cardiomyocytes. By contrast, *Mycn* demonstrated a unique and potent ability to drive both S-phase re-entry and mitotic progression in postnatal cardiomyocytes, as evidenced by significantly increased BrdU incorporation and pH3-positive nuclei (**Figures 3A**, **B**). These findings are consistent with its developmental role, as *Mycn* is essential for cardiomyocyte proliferation during cardiac embryogenesis and regeneration. Genetic ablation of *Mycn* during embryonic development results in myocardial hypoplasia via downregulation of key cell cycle regulators such as *Ccnd1* and *Ccnd2*, whereas *Myc* is dispensable in this context.^14,35^ Moreover, the Hedgehog–Gli1– Mycn signaling axis has been implicated in cardiac regeneration in both zebrafish and neonatal mice^36^. Our results extend these developmental insights into the postnatal heart, demonstrating that Mycn reactivates robust cell cycle activity in postnatal cardiomyocytes, evidenced by both S and M phase progression. Taken together, our data provide compelling evidence that *Mycn*, unlike its paralogs *Myc* and *Mycl*, possesses a non-redundant and unique capacity to drive cell cycle reactivation in cardiomyocytes, even in the postnatal heart.

### Potential Mechanisms of Mycn-mediated Cardioprotection

In this study, we demonstrated that *Mycn*-expressing postnatal hearts exhibit significantly preserved systolic function following MI—which we associate with reduced infarct area and increased capillary density in the peri-infarct region (**Figures 5B, C**). Cardiomyocyte-intrinsic changes alone are unlikely to account for the full extent of beneficial post-MI tissue remodeling observed in these *Mycn*-expressing hearts. While detailed mechanisms remain for future investigation, our transcriptomic analysis revealed that *Mycn* robustly upregulated a suite of genes associated with paracrine signalling, extracellular matrix remodeling, and stromal activation—many of which are enriched in neonatal cardiac environments, including *Fstl1*, *Timp1*, *Igf1*, *Postn*, *Ptn*, *Pf4*, *Col5a2*, *Col3a1*, and *Lum*—known to enhance angiogenesis, suppress inflammation, or promote cardiomyocyte survival under ischemic stress.^26–32^ For instance, FSTL1 is a known secreted factor that promotes angiogenesis and cardiomyocyte survival after MI;^26,27^ *Timp1* stabilizes the extracellular matrix and prevents adverse remodeling, and its deletion exacerbates cardiac dysfunction after myocardial injury;^28,29^ IGF1 enhances cardiomyocyte survival via the PI3K/Akt pathway and supports cardiomyocyte metabolic adaptation during ischemic stress.^30,31^ These factors secreted from *Mycn*-expressing cardiomyocytes may collectively contribute to constructing a cardioprotective microenvironment enhancing tissue resilience against irreversible ischemic injury.

In addition, we found that *Mycn*-expressing murine hearts exhibited increased numbers of Hsp47-positive fibroblasts and elevated collagen production, as shown by Picro-Sirius Red staining (**Figures 4 A-D**). Traditionally, fibroblast activation and collagen deposition are hallmarks of adverse remodeling and cardiac fibrosis. ^22,37,38^ However, emerging studies have identified functionally distinct fibroblast subtypes—including pro-angiogenic, immunomodulatory, and extracellular-matrix-stabilizing pheotypes— each responsive to paracrine cues from neighboring cardiomyocytes; notably, Hsp47 itself is essential for collagen maturation and is essential to prevent ventricular rupture during acute phases of cardiac injury. ^39–42^ This study showed absence of cardiac dysfunction at up to eight weeks, and preserved ejection fraction after MI in *Mycn*-expressing hearts, supporting the notion that rather than promoting pathological fibrosis, *Mycn*-induced fibroblasts may belong to reparative or protective subtypes.

### Correlation between Cell Cycle Activation and Cardioprotection after MI

Although direct mechanisms remain uninvestigated, the correlation between Mycn induced robust cell cycle re-entry and cardioprotection following MI observed in our study aligns with Bradley and colleagues’ recent findings—that existence of cycling cardiomyocytes correlates to cardioprotection from ischemic injury^9^. One potential explanation is that cell cycle activity serves as a surrogate marker of a broader phenotypic shift toward a fetal-like state of plasticity, characterized by altered gene expression profiles that confer injury resistance. To reinforce this idea, many of the Mycn-induced genes already discussed are abundantly expressed in naturally regenerative neonatal cardiomyocytes (**Supplemental Figure 1A**). Furthermore, in regenerative cardiac models (via YAP activation or transient OSKM induction), cardiomyocytes acquire fetal-like transcriptional signatures, acquiring enhanced stress tolerance and regenerative capacity.^43,44^—our data show that more than half of Mycn-induced genes overlapped with those highly expressed in neonatal cardiomyocytes (**Supplemental Figure 1B**), supporting the concept of fetal-like phenotype shift in postnatal cardiomyocytes. Thus, the correlation between cardiomyocyte cell cycle activity and cardiac protection may not reflect a direct causal relationship, but rather a phenotypic shift toward a fetal-like stress-adaptive state; this altered state may orchestrate cardioprotection through paracrine signaling and modulation of fibroblast phenotypes, ultimately constructing a supportive cardiac microenvironment.

## Conclusion

In summary, this study establishes *Mycn* as a functionally distinct member of the Myc family in the postnatal murine heart. *Mycn* robustly activates cardiomyocyte cell cycle re-entry, and confers significant cardioprotection following MI. This protective effect is likely mediated, at least in part, by reactivation of a fetal-like cardiomyocyte state, which coordinates stress-adaptive responses across cardiomyocytes and stromal cells alike. Our findings enhance the understanding of cardiac plasticity and support the therapeutic potential of *Mycn* or its downstream effectors in promoting cardiac repair in hearts under ischemic stress.

## Acknowledgments

We would like to thank Asami Kawamura for AAV production, Yoshitaka Tateishi for reagent preparation, members of the Institute of Biomedical Research at Sapporo-Higashi Tokushukai Hospital for their assistance with RNA-seq data analysis, and Asahikawa Medical University for providing Independent Research Support Grant for Fundamental Scientific Studies to AH.

## Abbreviations

AAV: adeno-associated virus
BrdU: 5-bromo-2′-deoxyuridine
cTnT: cardiac troponin T
EF: ejection fraction
FFPE: formalin-fixed paraffin-embedded
GFP: green fluorescent protein
GO: Gene Ontology
HE: hematoxylin and eosin
LAD: left anterior descending (artery)
MI: myocardial infarction
PSR: picrosirius red
pH3: phospho-histone H3
qPCR: quantitative polymerase chain reaction
RNA-seq: RNA sequencing
RT-PCR: reverse transcription polymerase chain reaction
WGA: wheat germ agglutinin

---

**Supplemental Figure 1.**
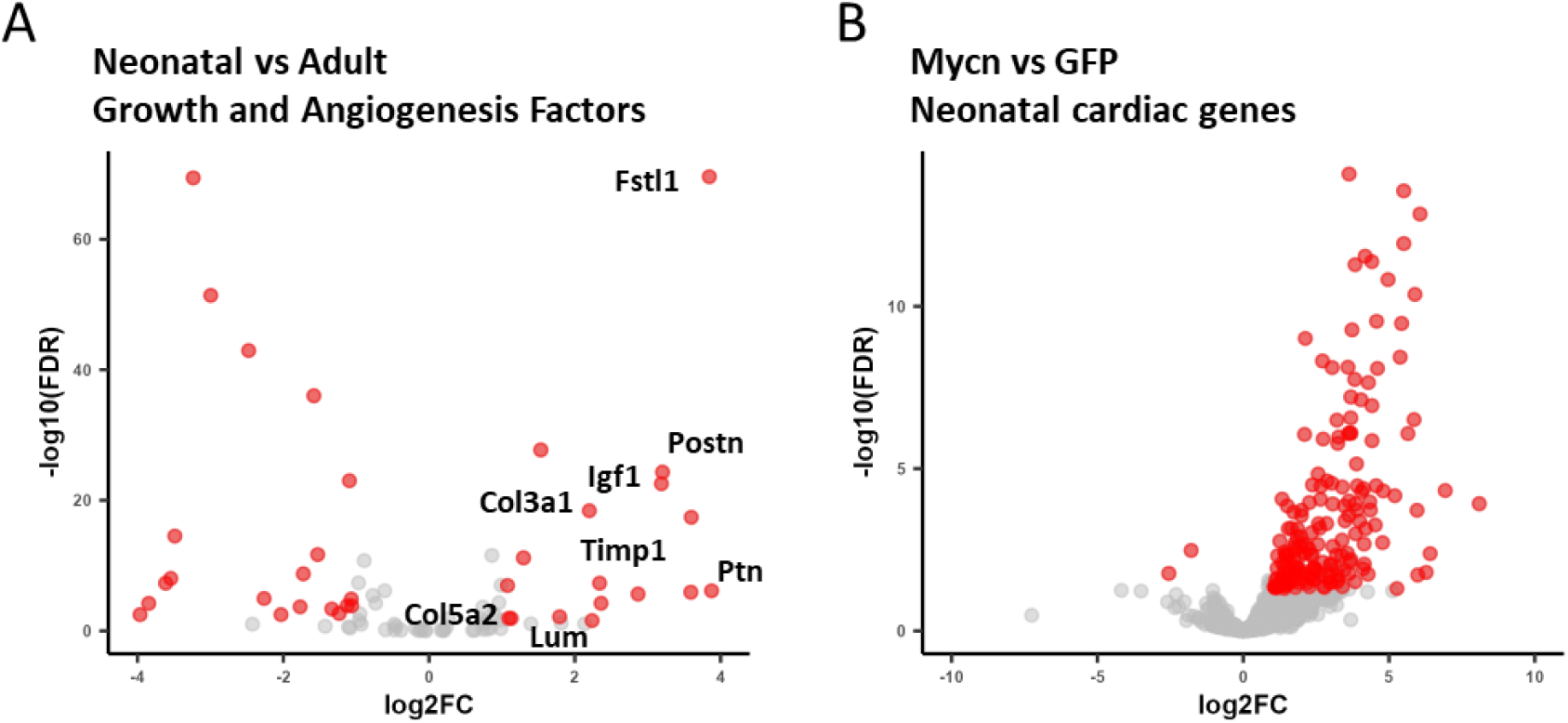
**A. *Mycn*-induced growth and angiogenic factors are highly expressed in neonatal cardiomyocytes:** To assess the developmental expression patterns of Mycn-induced growth and angiogenic factors, a previously published RNA-seq dataset (NCBI Accession: SRP033386) comparing neonatal (P1) and adult cardiomyocytes was reanalyzed using the same pipeline described in the Methods section. Genes associated with the Gene Ontology term “growth factor activity” (GO:0008083) or included in a curated angiogenesis gene set were visualized in a volcano plot. Mycn-induced growth and angiogenic factors were specifically labeled, revealing that a substantial number of these growth and angiogenesis-related factors are also highly expressed in neonatal cardiomyocytes. Genes with a false discovery rate (FDR) < 0.05 and a fold change (FC) ≥ 2 (neonatal cardiomyocytes vs adult cardiomyocytes) are shown in red. **B. Upregulation of neonatal genes in *Mycn*-expressing cardiomyocytes:** Using the same RNA-seq dataset as in Supplemental Figure 1A, neonatal cardiac genes were defined as those significantly upregulated in P1 cardiomyocytes relative to adult cardiomyocytes (false discovery rate (FDR) < 0.05 and absolute fold change (FC) ≥ 2), and compiled into a neonatal gene list. Differential expression analysis was then performed between *Mycn*- and *GFP*-expressing cardiomyocytes to assess the expression of these neonatal genes. The results were visualized as a volcano plot. Of the 367 genes upregulated in the Mycn group, 188 genes (51%) overlapped with the neonatal gene list, a phenotypic shift toward a neonatal-like cardiomyocyte state.

## Supplemental file 1

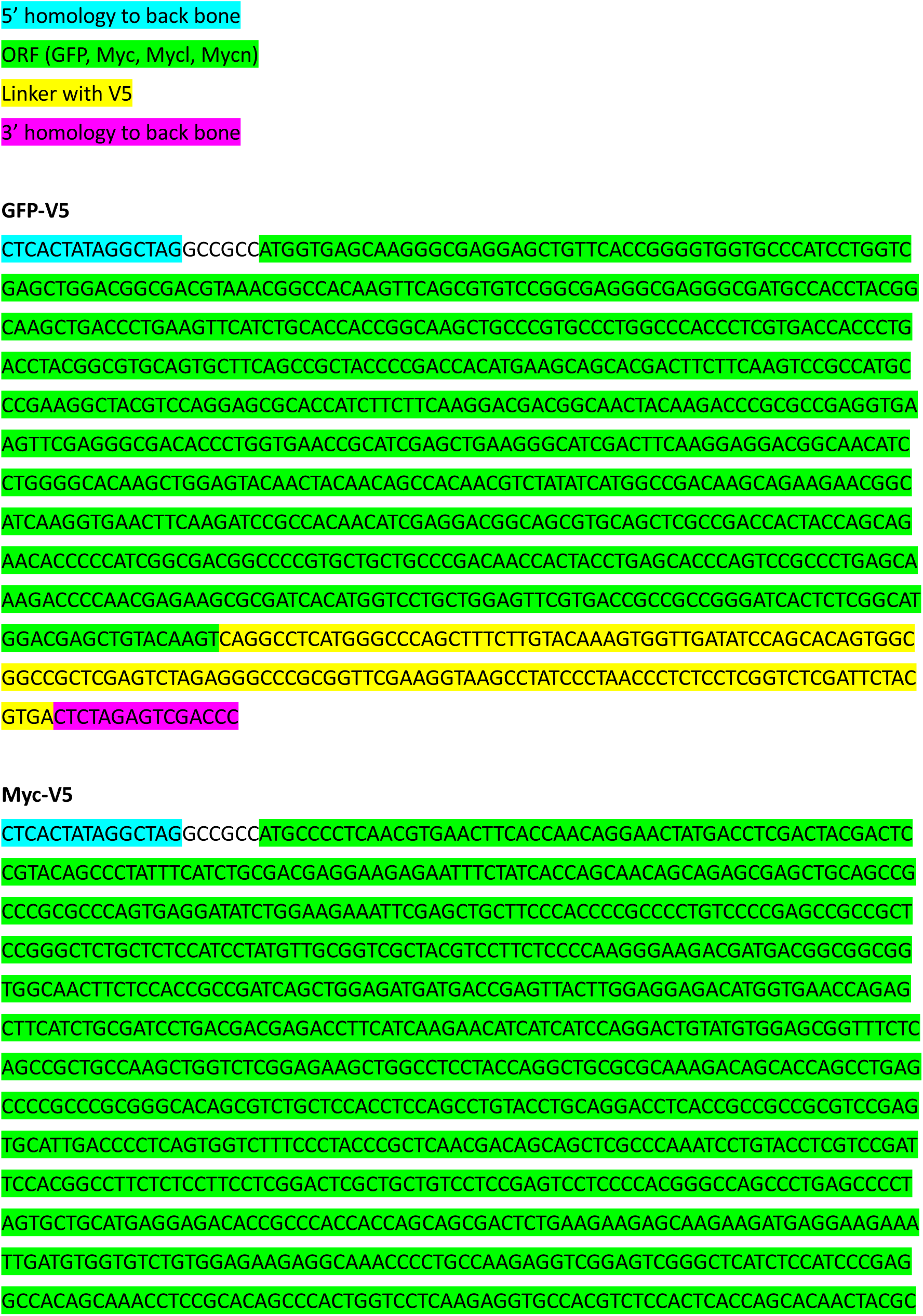
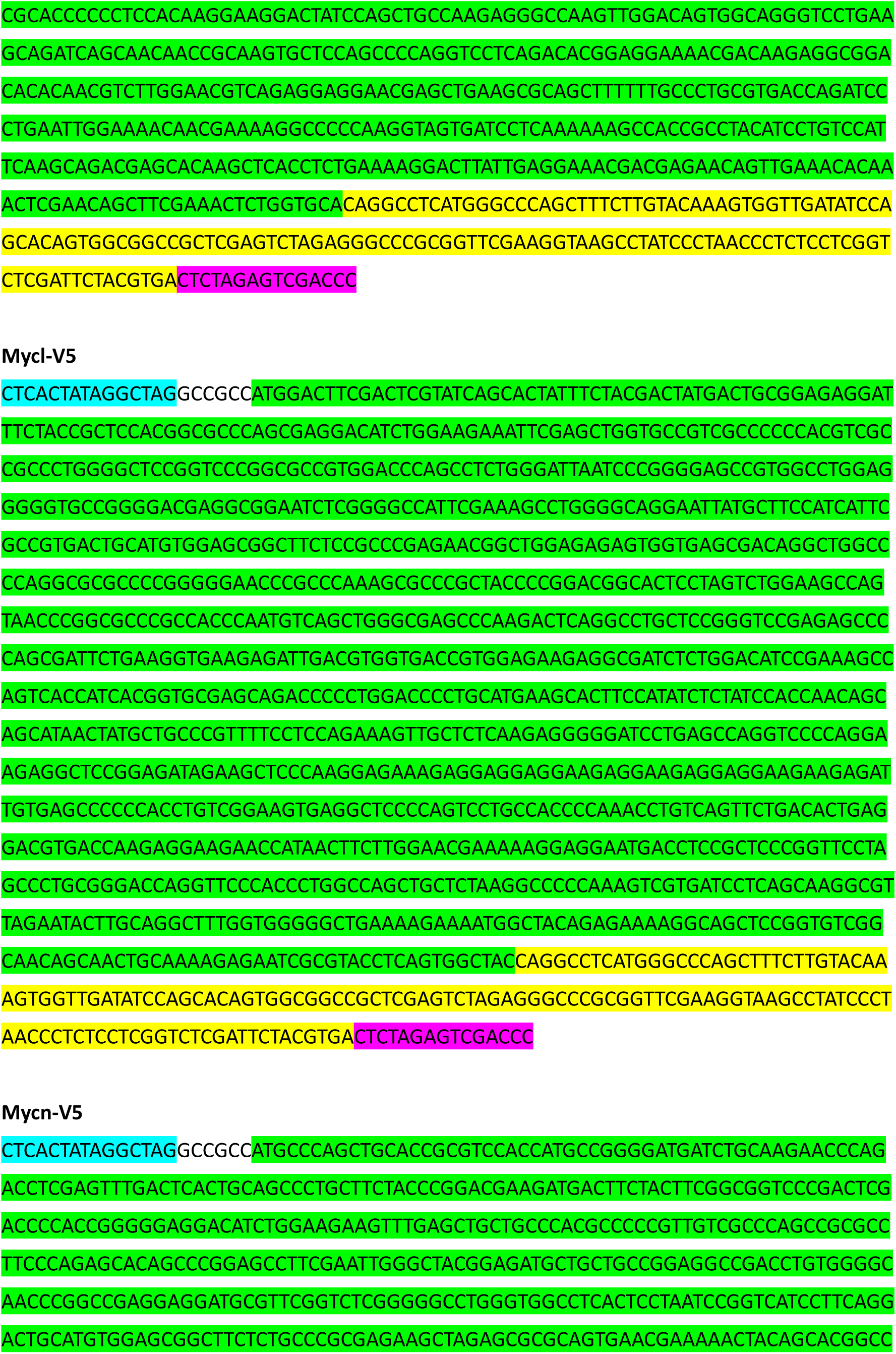
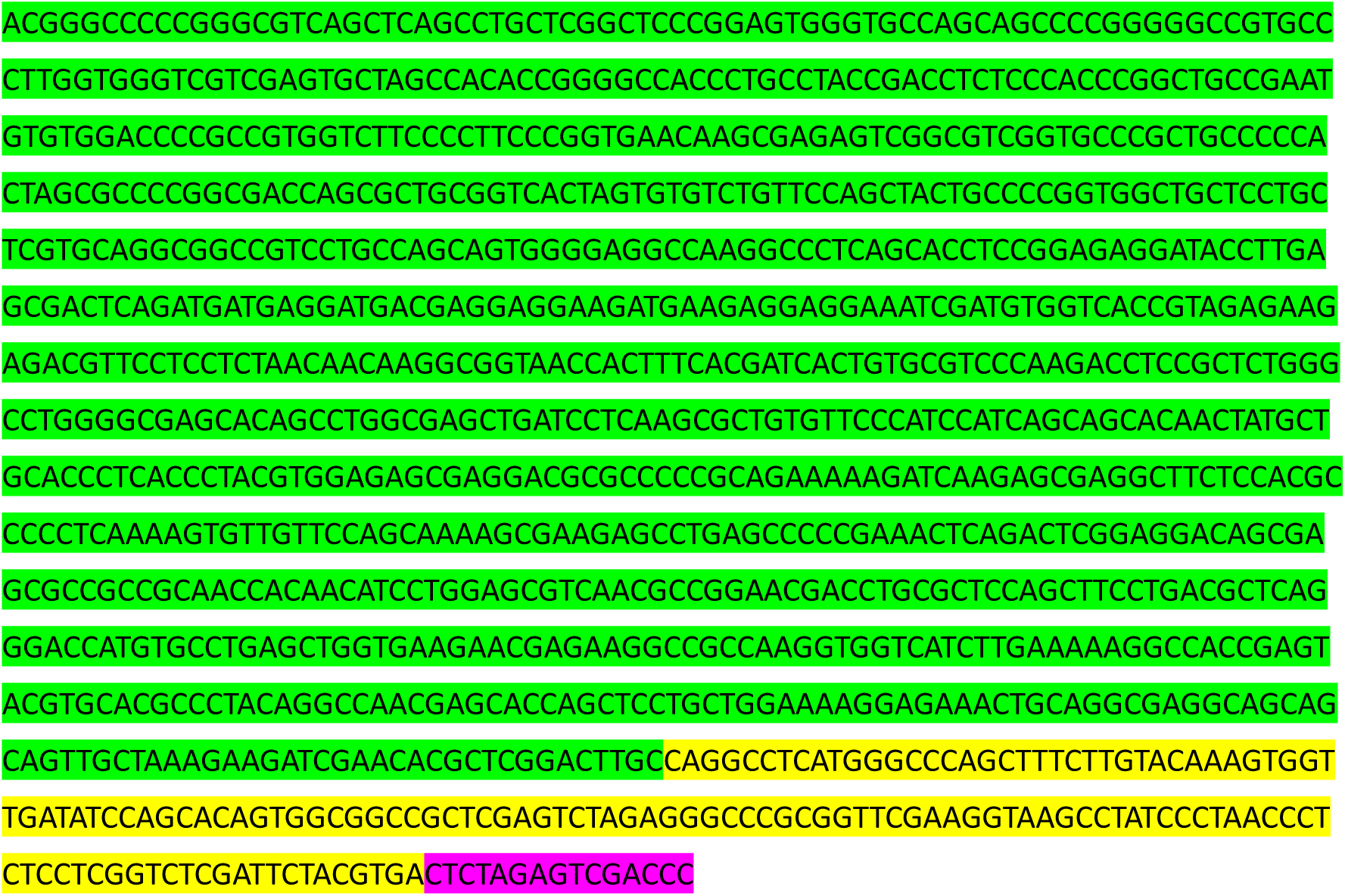

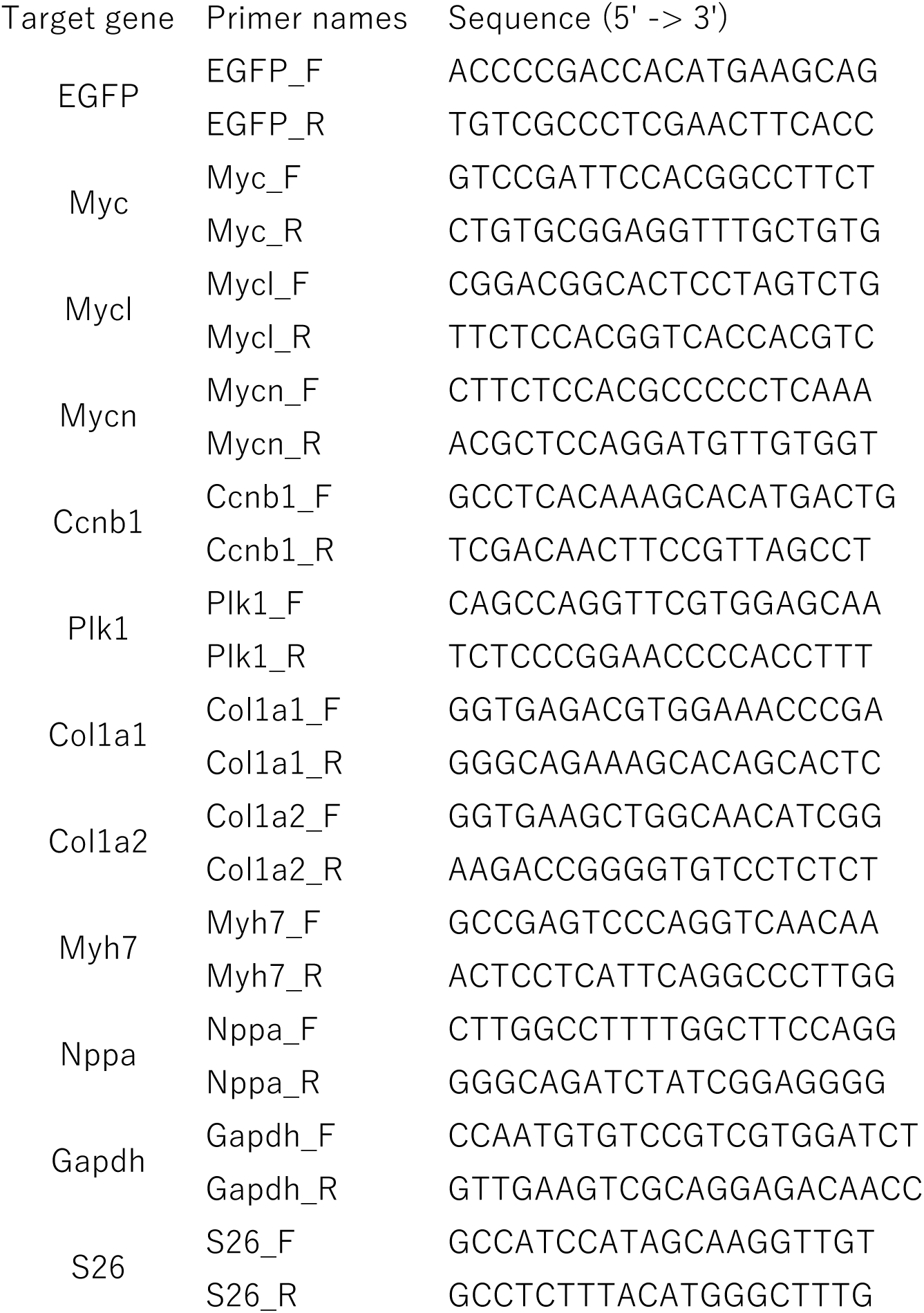

